# Monoacylglycerol acyltransferase maintains ionotropic receptor expression for cool temperature sensing and avoidance in *Drosophila*

**DOI:** 10.1101/2024.08.02.606314

**Authors:** Xiangmei Deng, Takuto Suito, Makoto Tominaga, Takaaki Sokabe

## Abstract

Sensory inputs of temperature dynamics in the environment are essential for appropriate physiological outputs. The responsiveness of sensory neurons is maintained by functional thermosensor expression. However, the mechanism by which their expression is regulated is unclear. In this study, we identified a monoacylglycerol acyltransferase-coding gene named *bishu-1* that contributes to maintaining the responsiveness of cool temperature sensing neurons in *Drosophila*. *bishu-1* mutation led to abnormal thermal avoidance in a cool temperature range. Cooling-induced responses in dorsal organ cool cells were weakened by the absence of *bishu-1*, and this was associated with reduced transcription of the ionotropic receptors *IR25a* and *IR21a* through the transcription factor *broad*. Our findings unveil a novel link between lipid metabolism and thermosensor function, thus providing new insights into mechanisms underlying the appropriate maintenance of sensory inputs.

## Introduction

Environmental temperature is a critical parameter for all animals as they seek optimal habitats for survival and successful reproduction. To monitor surrounding temperatures, a wide range of species have evolved temperature sensing mechanisms involving thermosensitive molecules including transient receptor potential (TRP) channels (1), other types of ion channels (2, 3), and G protein-coupled receptors (GPCRs) (4–7). Thermal sensation is particularly important for ectotherms such as insects because of their small body size and large surface-to-volume ratio, which lead to rapid core temperature fluctuation. Thus, their physiological functions, including long-term survival (8), reproduction (9), and diverse behavioral activities (10), are profoundly impacted by ambient temperature. To mitigate core temperature fluctuation, *Drosophila* has evolved a delicate thermosensory system that discriminates temperature changes within a milli-degree per second (11). This accurate thermosensation relies on the precise maintenance of thermosensor functions. In addition to GPCRs and TRP channels (4, 6, 12), *Drosophila* larvae and adults use subsets of unique receptor families identified in invertebrates, such as ionotropic receptors (Irs) and a gustatory receptor for thermosensation (13–17). However, the activation and regulatory mechanisms of many of these thermosensors remain to be clarified.

Lipids are important players in TRP channel regulation and sensory responses. For example, fatty acids serve as key signaling molecules that modulate TRP channels in multiple species (18). In the aspect of thermosensory and regulatory behaviors in *Drosophila*, lipid products of phospholipase Cβ (PLC) signaling cascade and other membrane lipids have been reported; the thermal preference of larvae is influenced by desaturation and oxidization of membrane lipids (19–21). In adults, cold temperature activates bitter taste neurons expressing rhodopsin and PLC, which is essential for cold temperature dependent feeding suppression (22). Low ambient temperature facilitates unsaturated fatty acid intake and egg laying on unsaturated fatty acid-containing food in adults (23), suggesting the importance of saturation level of membrane lipids in maintenance of biological functions. PLC signaling cascade is also involved in other sensory processes of *Drosophila.* In phototransduction, products of PLC including diacylglycerol (DAG) and monoacylglycerol (MAG) activate TRP and TRPL channels (24–26). The same or related PLC products possibly regulate TRPA1 in taste neurons in response to bitter compounds (27). These facts suggest that diacylglycerol and relevant metabolites in PLC and other signaling cascades are key components for receptor and sensory functions. However, the regulatory mechanisms and physiological importance of associated lipid enzymes in thermosensory processes have not been thoroughly clarified.

Transcriptional regulation of sensors by lipids is one possible mechanism to maintain receptor function. In support of this, accumulated evidence links lipids and transcription. For example, polyunsaturated fatty acids function as ligands of nuclear receptors such as peroxisome proliferator-activated receptor (PPAR) and the transcription factor sterol regulatory element binding protein (SREBP) in multiple tissues in mammals (28, 29). DAG is an activator of protein kinase C (PKC), which is coupled with the activation of multiple transcription factors such as NFқB, AP-1 (30), signal transducers and activators of transcription 3 (Stat3) (31), and c-Rel (32). Additionally, a recent report suggested that the activity of the Max-like protein X (MLX) family of transcription factors is suppressed through binding to lipid droplets, which are formed by DAG and acylglycerol acyltransferases (33). However, whether fatty acids and DAG affect transcription in neurons remains unknown.

In this study, we conducted thermotaxis screenings using *Drosophila* larvae and identified two DAG synthesis genes, namely *bishu-1* and *bishu-2*. The name “*bishu*” given to the gene is a Chinese word for “summering,” describing the escaping behavior of larvae from heat. *bishu-1* and *bishu-2* mutation led to defects in cool temperature avoidance. These genes formed a cluster on the genome, and their amino acid sequences were close to those of the human diacylglycerol acyltransferase 2 (DGAT2) family. *bishu-1* and *bishu-2* were distinctly expressed in multiple sensory neurons to regulate thermal preference, and *bishu-1* particularly functioned in cool temperature-sensitive dorsal organ cool cells (DOCCs) expressing a set of thermosensors *IR21a*, *IR25a*, and *IR93a*. The abnormal cool temperature avoidance in *bishu-1^KO^* was sufficiently rescued by human monoacylglycerol acyltransferase (MGAT) genes, suggesting the importance of their conserved function as acyltransferases in thermosensation. *In vivo* GcaMP imaging revealed that the response of DOCCs to cool temperature was diminished in the *bishu-1^KO^*. The reduced responses of *bishu-1 ^KO^* DOCCs was attributed to the downregulation of *IR21a* and *IR25a* mediated by the transcription factor *broad* (*br*). These results unveil a novel mechanism by which lipid metabolism regulates receptor function in sensory neurons through maintaining gene expression.

## Results

### Putative acylglycerol acyltransferases are involved in cool temperature avoidance

We conducted a thermal preference assay to screen *Drosophila* strains carrying P-element insertions in genes involved in metabolism of DAG and fatty acids (unpublished) and identified the putative acylglycerol acyltransferase *CG1942*/*bishu-1*. *bishu-1* formed a tandem cluster on the genome with two structurally related genes, namely *CG1941* and *CG1946/bishu-2* (Fig. S1*A*). Phylogenetic analysis with amino acid sequences from human DGAT1 and DGAT2 families indicated that proteins encoded by *CG1941*, *bishu-1*, and *bishu-2* have a closer evolutionary relationship with human DGAT2 family (Fig. 1*A*). This family contains MGATs, DGATs, and acyl-CoA wax alcohol acyltransferase (AWATs). Among all, Bishu-1 displayed higher amino acid identity with MGATs (MGAT2: 39.52%, MGAT3: 38.12%, MGAT1: 35.52%) than other members (DGAT2: 30.67%, AWAT1 35.06%). Because the molecular function of human MGATs are known to produce DAG using acyl-CoA and MAG as substrates (34–36), we predicted *bishu-1* as a DAG synthesis enzyme-coding gene. To study the effect of loss of function of each gene on thermal preference, we generated knockout lines of *CG1941*, *bishu-1*, and *bishu-2* using the CRISPR-Cas9 technique. *CG1941^KO^* was generated by introducing a deletion in the first exon to induce a frameshift (Fig. S1*B*, top), whereas *bishu-1^KO^* and *bishu-2^KO^* were generated by inserting a *DsRed* reporter downstream of the start codon to interrupt their expression (Fig. S1*B*, middle and bottom). The mRNA level of the target gene was abolished in each allele, indicating they were null mutants (Fig. S1*C*). The mRNA level of a neighboring gene in each knockout was reduced by approximately 50% despite the lack of a mutation in its coding sequence, suggesting that the expression of each gene is maintained reciprocally (Fig. S1*C*).

**Fig. 1.**
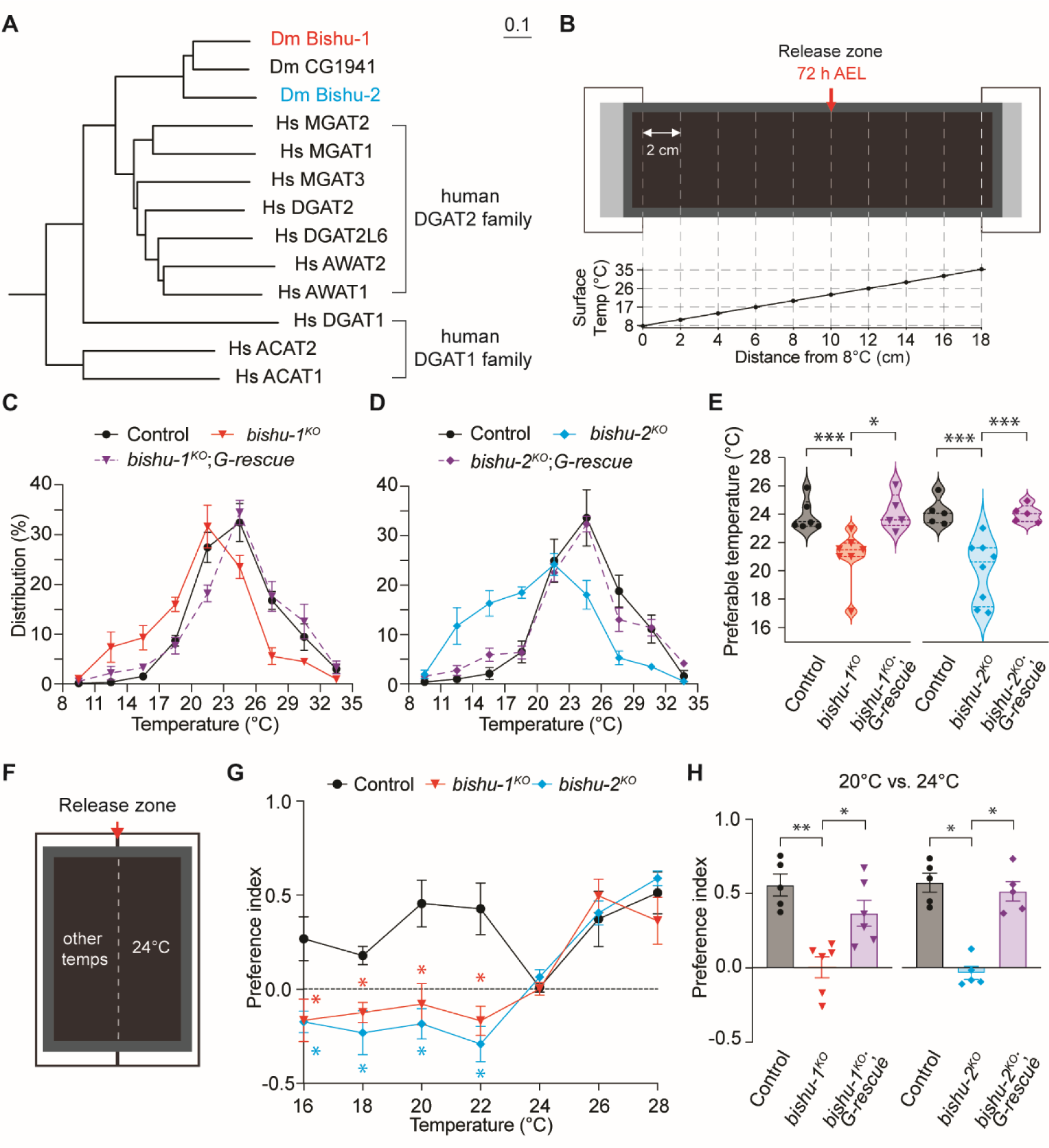
*bishu-1* and *bishu-2* were involved in cool temperature avoidance. (A) Phylogenetic tree of the *Drosophila* proteins (Dm) CG1941, Bishu-1 and Bishu-2 and the human proteins (Hs) from the DGAT1 family (DGAT1, ACAT1, and ACAT2) and DGAT2 family (DGAT2, DGAT2L6, MGAT1, MGAT2, MGAT3, AWAT1, and AWAT2). The scale bar indicates the p-Distance. (B) Schematic diagram of the thermal gradient assay. An aluminum testing plate was coasted with 2% agarose and placed on two separated aluminum blocks (white boxes), each of which was set up with a distinct temperature using circulating water bath. Early third instar larvae (72 h AEL) were released at 23°C (red arrow). After releasing larvae, a glass lid was placed on the testing plate during the assay. The actual temperatures measured on the testing plate with the glass lid are presented. Data are presented as the mean ± SD (N = 3). Error bars are not visible as the variance is smaller than the marker size. (C–E) Distribution of early third instar larvae (72 h AEL) of each genotype on an 8°C–35°C linear gradient (C and D) and the preferable temperature (E). The *G-rescue* denotes the wild-type of a genomic transgene including *CG1941, bishu-1*, and *bishu-2*. (C) *w^1118^* (control), *bishu-1^KO^*, and *bishu-1^KO^*;*G-rescue* (N= 5–7). (D) Control, *bishu-2^KO^*, and *bishu-2^KO^*;*G-rescue* (N = 5–8). (E) The preferable temperature indicates the median staying temperatures, which was calculated using the distribution of larvae in C (left) and D (right). (F) Schematic diagram of the thermal two-way choice assay. An aluminum testing plate was coated with 2% agarose and placed on two adjacent aluminum blocks. Larvae were released at the center zone (red arrow). (G) Preference indices (PIs) of two-way choice assays between 24°C versus other temperatures. Larvae of control, *bishu-1^KO^*, and *bishu-2^KO^* (N = 4–7). (H) PIs of two-way choice assays in a 20°C versus 24°C condition. Control, *bishu-1^KO^*, *bishu-1^KO^*;*G-rescue* (left); *bishu-2^KO^* and *bishu-2^KO^*;*G-rescue* (right, N = 5–6). The data are presented as the mean ± SEM except for (B). Each point represents a biological replicate. *\*P* < 0.05, *\*\*P* < 0.01, and *\*\*\*P* < 0.001 by one-way ANOVA with Tukey’s test (E, and H) or Dunnett’s test (G). NS denotes not significant.

We tested the thermal preference of these knockouts on a linear temperature gradient of 8°C–35°C (Fig. 1*B*), covering optimal and noxious temperature ranges. We released cohort of early third instar larvae [72 h after egg laying (AEL)] at 23°C and allowed them to explore the thermal landscape on the plate under a dim red light (>600 nm). Control larvae most accumulated in the 23°C–26°C zone (32.47% ± 3.84%, Fig. 1*C*; 33.60% ± 5.69%, Fig. 1*D*). Their preferable temperature, which is the median staying temperature of larvae on the gradient, was approximately 24°C [24.18°C ± 0.45°C (left) and 24.14°C ± 0.42°C (right), Fig. 1*E*]. The thermal preference of *CG1941^KO^* was comparable to that of the control, with the largest population in the 23°C–26°C zone (38.53% ± 2.70%, Fig. S1*D*) and the preferable temperature was 23.35°C ± 0.27°C. By contrast, *bishu-1^KO^* and *bishu-2^KO^* tended to distribute in a cooler range (11°C–20°C) with the largest population in the 20°C–23°C zone (31.65% ± 4.30% in *bishu-1^KO^* and 24.17% ± 2.23% in *bishu-2^KO^*, Fig. 1 *C* and *D*). The preferable temperature of *bishu-1^KO^* and *bishu-2^KO^* were significantly reduced to 21.30°C ± 0.70°C and 19.96°C ± 0.80°C, respectively (Fig. 1*E*). On the other hand, the control and all three knockouts consistently avoided the lowest temperature zone (8°C–11°C). The defect in cool temperature avoidance by *bishu-1^KO^* and *bishu-2^KO^* was rescued by introducing a genomic insertion containing the wild-type alleles of *CG1941*, *bishu-1*, and *bishu-2* (Fig. 1 *C–E*). These results suggested that the putative acylglycerol acyltransferases are specifically involved in the avoidance of innocuous cool temperatures.

Because *Drosophila melanogaster* larvae display a development-dependent switch in thermal preference from warm to cool temperatures (6, 37), we evaluated the developmental rate of the control and the knockouts. The proportion of third instar stage at 74 h AEL and the pupation timing did not differ among the genotypes (Fig. S1 *E* and *F*). In addition, we observed that *bishu-1^KO^* and *bishu-2^KO^* displayed a less visible shift in distribution to cooler ranges on the thermal gradient both at the second (48 h AEL, Fig. S1*G*) and late third instar stages (120 h AEL, Fig. S1*H*) compared to the early third instar stage (72 h AEL, Fig. 1*C* and *D*).

To further confirm that cool temperature avoidance depends on *bishu-1* and *bishu-2*, we conducted a thermal two-way choice assay (Fig. 1*F*). Based on the preferable temperature of the control on the thermal gradient (Fig. 1*E*), we set 24°C as a reference against the test temperatures. We released larvae at a border of 24°C and test temperature between 16 and 28°C with a 2°C interval and allowed them to choose two temperature zones. We counted the number of larvae on each side and calculated the preference index (PI). We observed that control and *CG1941^KO^* larvae preferred 24°C to cooler or warmer testing temperatures (Fig. S1*I*), which was consistent with the gradient assay results. *bishu-1^KO^* and *bishu-2^KO^* avoided temperatures exceeding 24°C similarly as the control, but they more frequently accumulated at temperatures lower than 22°C (Fig. 1*G*). This inability to avoid cool temperatures in *bishu-1^KO^* and *bishu-2^KO^* was rescued by introducing their wild-type alleles (Fig. 1*H*). These results strengthen the idea that *bishu-1* and *bishu-2* specifically contribute to cool temperature avoidance. Because a 20°C versus 24°C condition yielded the highest PI in the control, we performed subsequent two-way choice assays under this condition.

### *bishu-1* primarily functions in DOCCs to mediate cool temperature avoidance

To explore the responsible cells for *bishu-1*– and *bishu-2*–dependent thermotaxis, we performed RNAi-mediated knockdown of each gene using the GAL4-UAS expression system. A pan-neuronal driver (*elav-GAL4*)–induced knockdown of *bishu-1* or *bishu-2* resulted in a significant reduction in PI in the 20°C versus 24°C condition (Fig. S2*A*), suggesting that these genes function in neurons to discriminate temperatures. Next, we suppressed *bishu-1* or *bishu-2* in sensory neurons expressing thermosensors involved in cool temperature sensation including TRPL, Inactive (Iav), and Irs (12, 15, 37). Knockdown of *bishu-1* or *bishu-2* in TRPL-expressing neurons did not affect PI (Fig. S2*B*), whereas PI was significantly reduced when *bishu-1* or *bishu-2* was suppressed under the control of *iav-GAL4* (Fig. S2*C*). However, the heterozygous *GAL4* control (*iav-GAL4/+*) displayed variable Pis with a lower average value than other controls, which complicated interpretation of the result.

Previous studies reported that *IR25a*, *IR93a*, and *IR21a* are co-expressed in DOCCs to mediate cool temperature avoidance (15, 37). When we suppressed *bishu-1* or *bishu-2* using *IR25a-GAL4*, obvious impairments in temperature discrimination were observed between 20°C and 24°C (Fig. 2 *A* and *B*), suggesting possible functions of these genes in DOCCs. Consistent with this idea, the defect in cool temperature avoidance in *bishu-1^KO^* was rescued by overexpressing *bishu-1* cDNA using *IR25a-GAL4* (Fig. 2*C*). By contrast, overexpressing *bishu-2* using *IR25a-GAL4* failed to rescue the phenotype in *bishu-2^KO^* (Fig. 2*D*). To further confirm the functions of *bishu-1* and *bishu-2*, we suppressed each gene using DOCC-specific *IR21a-GAL4* or *R11F02-GAL4* (11, 15). Defects in cool temperature avoidance were observed upon *bishu-1* knockdown, but not *bishu-2* knockdown, in DOCCs (Fig. 2 E and F). *R11F02-GAL4*–induced overexpression of *bishu-1* in the *bishu-1^KO^* background sufficiently recovered the preference for 24°C (Fig. 2*G*). Moreover, considering the sequence similarity between *bishu-1* and the human DGAT2 family, we overexpressed three human genes from this family in *bishu-1^KO^* and observed sufficient compensation for the *bishu-1^KO^* phenotype by two MGATs, but not by DGAT2 (Fig. 2*H*). Taken together, these results strongly suggested that *bishu-1* and *bishu-2* function in multiple but distinct tissues to mediate cool temperature avoidance. Specifically, *bishu-1* emerged as a primary contributor in DOCCs with possible MGAT activity.

**Fig. 2.**
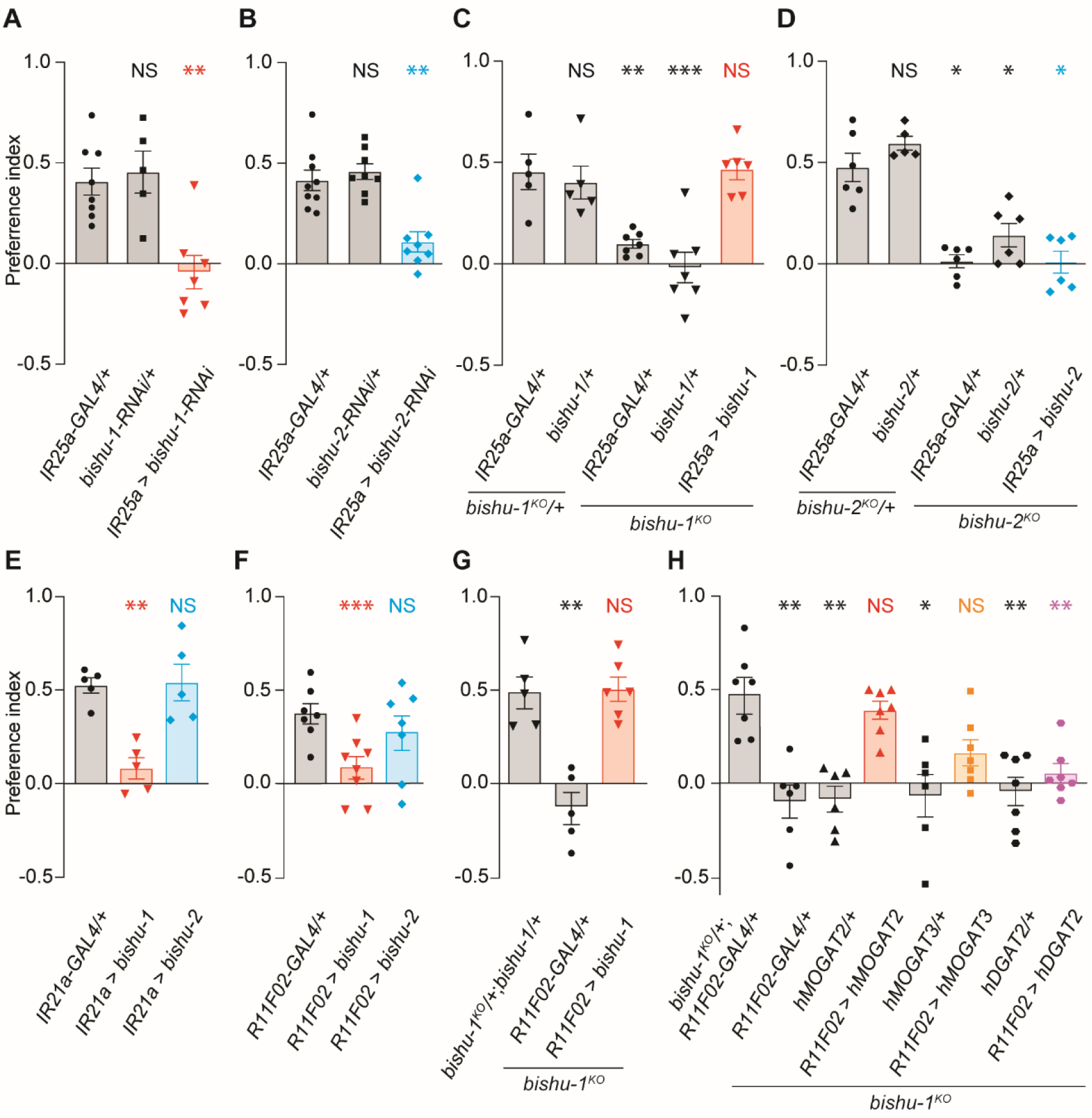
*bishu-1* functioned in cool temperature avoidance in DOCCs. Preference indices (PIs) of two-way choice assays in a 20°C versus 24°C condition with the indicated genotypes. (A and B) Knockdown of *bishu-1* (A) or *bishu-2* (B) in *IR25a*-expressing neurons (N = 5–9). (C and D) Rescue of *bishu-1^KO^* (C) or *bishu-2^KO^* (D) by introducing a wild-type transgene under the control of *IR25a-GAL4* (N = 5–7). (E and F) Knockdown of *bishu-1* or *bishu-2* in DOCCs using a specific driver *IR21a-GAL4* (E) or *R11F02-GAL4* (F, N = 5–8). (G) Rescue of *bishu-1^KO^* by overexpressing wild-type *bishu-1* in DOCCs using *R11F02-GAL4* (N = 5–6). (H) Thermal choice of *bishu-1^KO^* overexpressing human genes. Transgenes of human *MGAT2* (*UAS-hMGAT2*), *MGAT3* (*UAS-hMGAT3*), and *DGAT2* (*UAS-hDGAT2*) were expressed in *bishu-1^KO^* using *R11F02-GAL4* (N = 5–7). The data are presented as the mean ± SEM. \**P* < 0.05, \*\**P* < 0.01, and \*\*\**P* < 0.001 by one-way ANOVA with Dunnett’s test (A–C and E–H) or the Kruskal–Wallis test with Steel’s test (D). NS denotes not significant.

### The absence of *bishu-1* affects the cooling response of DOCCs

The functional contribution of *bishu-1* in DOCCs led us to examine whether the gene is expressed in *IR25a* and *IR21a* neurons. We generated a *bishu-1–*P2AQF2 fused line (Fig. S3*A*) and examined the *bishu-1* expression pattern using QUAS-mCherry as a reporter. In the anterior region of larvae, *IR25a* is expressed in temperature sensitive neurons including DOCCs and dorsal organ warm cells (DOWCs) (17). We observed that *bishu-1* expression largely overlapped with *IR25a* neurons (Fig. 3 *A*–*C*). Among them, three *bishu-1*–expressing neurons were DOCCs because they were co-labeled with *IR21a*-*GAL4* (Fig. 3*D–F* and S3*B*).

**Fig. 3.**
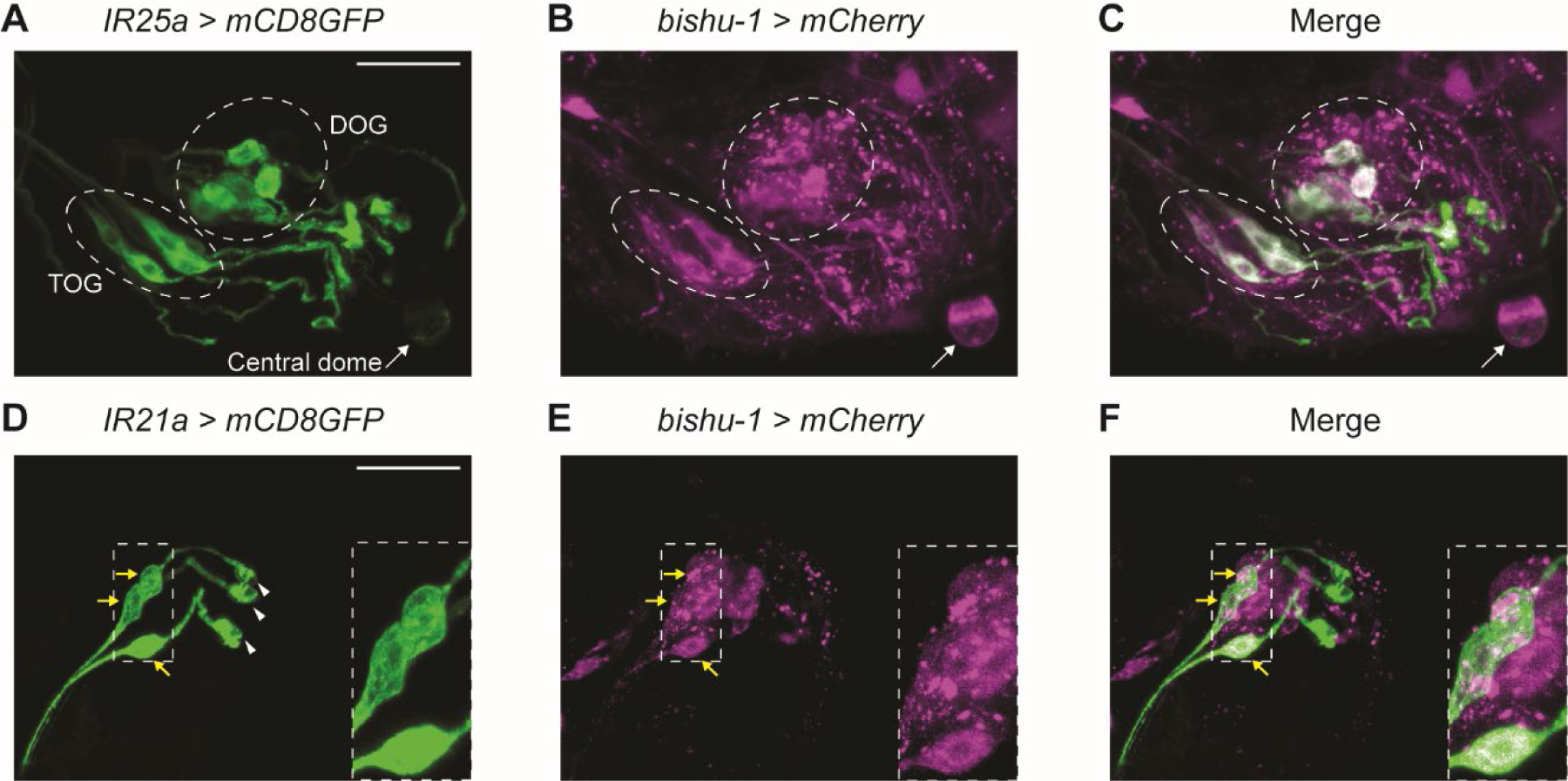
*bishu-1* was expressed in *IR25a*- and *IR21a*-expressing neurons. (A–C) Representative images of *bishu-1* and *IR25a* reporter expression in the anterior region of larvae. *IR25a-GAL4* (A, *IR25a-GAL4/+*;*UAS-40×GFP/+*), *bishu-1-P2AQF2* (B, *10×QUAS-6×mCherry/+*;*bishu-1-P2AQF2/+*), and the merged image (C). Dashed circles indicate the DOG and terminal organ ganglion (TOG), whereas the right arrow indicates the central dome, a perforated cuticular structure at the tip of the head (A–C). (D–F) Representative images of *bishu-1* and *IR21a* reporter expression in the anterior region of larvae. *IR21a-GAL4* (D, *IR21a-GAL4/+*;*UAS-40×GFP/+*), *bishu-1-P2AQF2* (E, *10×QUAS-6×mCherry/+*;*bishu-1-P2AQF2/+*), and the merged image (F). Yellow arrows and white arrowheads denote the cell bodies (D–F) and dendritic bulbs of DOCCs (D), respectively. The positions of the cell bodies of DOCCs (D–F) were highlighted as magnified insets in dashed boxes. In all images, the right side is the anterior end. Scale bars in A and D indicate 20 μm. The expression pattern was investigated in three or more individuals.

To explore the physiological role of *bishu-1* in DOCCs, we first examined the morphology of DOCCs in *bishu-1^KO^* and observed no obvious changes in the cell bodies and dendritic bulbs, a unique structure in DOCCs (11) (Fig. S4). Previous studies reported that cooling responses in DOCCs rely on *IR25a* and *IR21a*, and these responses are required for the avoidance of innocuous cool temperatures (15, 37). Because *bishu-1* was co-expressed with *IR25a* and *IR21a*, we performed *in vivo* Ca^2+^ imaging to evaluate the thermal responsiveness of DOCCs. We expressed *GcaMP8m* in DOCCs using *R11F02-GAL4* and subjected these neurons to temperature fluctuation between 18°C and 24°C. Both control and *bishu-1^KO^* DOCCs responded to temperature increases and decreases (Fig. 4 A–C). However, the cooling-induced Ca^2+^ increase was significantly reduced in *bishu-1^KO^* (Fig. 4 C–E). The reduction was evident in the cooling-induced peak responses and following sustained phases during cooling stimulation. The defect was partially restored by overexpressing *bishu-1* transcripts in the DOCCs using the *R11F02-GAL4* (Fig. 4 C–E). Taken together, these results indicate that the loss of *bishu-1* affects the cool responsiveness of DOCCs.

**Fig. 4.**
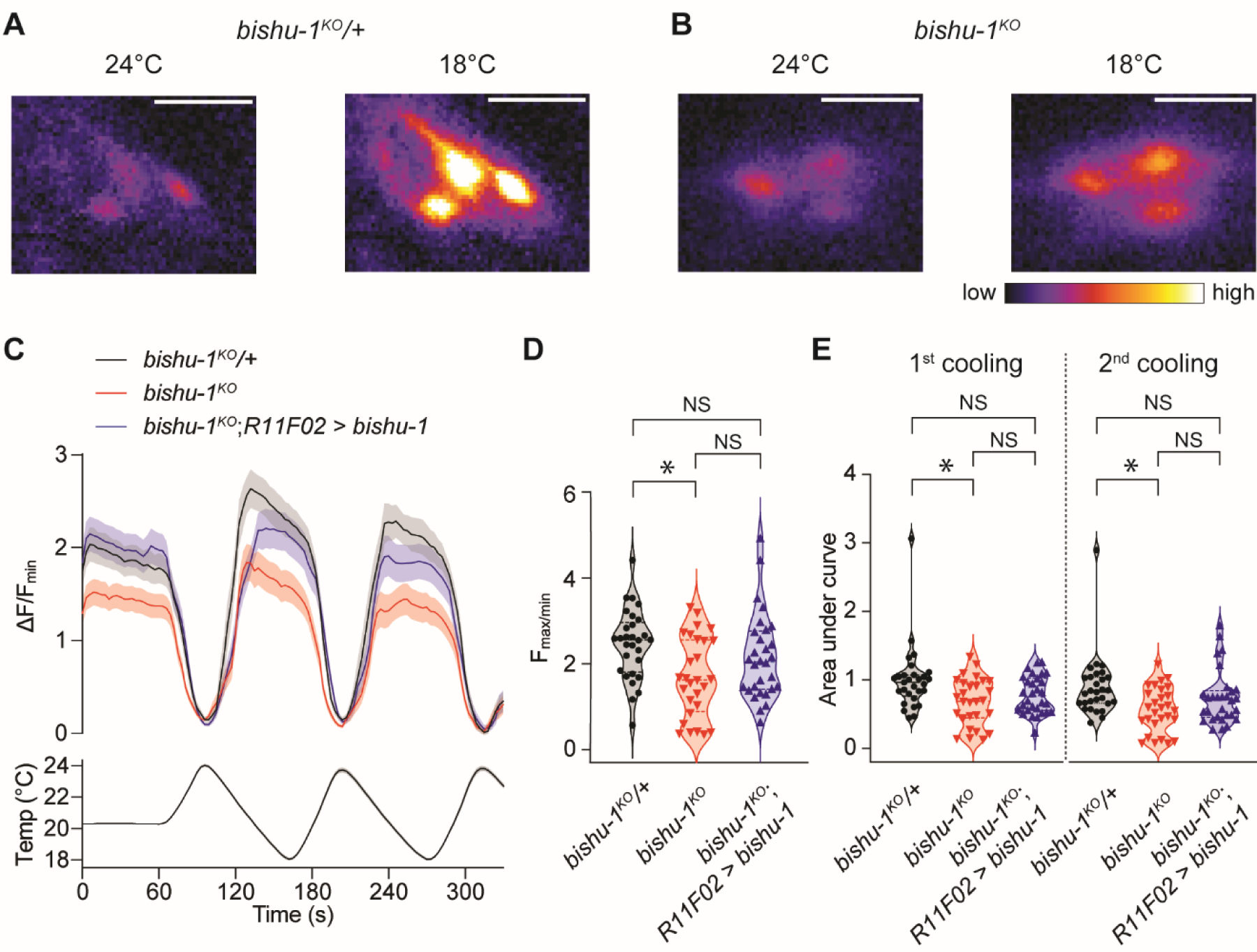
Loss of *bishu-1* caused the deterioration of cooling responses in DOCCs. (A and B) Representative GCaMP8 responses of DOCCs in the heterozygous control (*bishu-1^KO^*/+, A) and *bishu-1^KO^* (B) at 18°C or 24°C. GCaMP8 was expressed in DOCCs using *R11F02-GAL4*. Scale bars indicate 20 μm. (C) GCaMP responses (ΔF/F_min_) in *bishu-1^KO^*/+ (black), *bishu-1^KO^* (red), and a DOCC-specific rescued *bishu-1^KO^* (blue). F_min_ was taken between 60 s and 300 s. (D) Quantification of the averaged maximum Ca^2+^ responses of two cooling processes in (C). (E) Quantification of the area under the curve during the first (96–162 s) and second (204–270 s) cooling processes. All calculated results were normalized to the first cooling response of *bishu-1^KO^*/+. The data are presented as the mean ± SEM. *bishu-1^KO^*/+: N = 29 cells (from 12 animals); *bishu-1^KO^*: N = 28 (from 11 animals); *bishu-1^KO^*;*R11F02 > bishu-1*: N = 30 (from 13 animals). \**P* < 0.05 by the Kruskal–Wallis test with the Steel–Dwass test. NS denotes not significant.

### *bishu-1* regulates *IR25a* mRNA levels through the transcription factor *broad*

A previous study claimed that thermal preference was synchronized with expression changes in IR25a, IR21a and IR93a (37). Specifically, high or low expression of these receptors was accompanied by avoidance or acceptance of cool temperatures, respectively. Therefore, we examined the transcriptional levels of *IR25a*, *IR21a*, and *IR93a*, which function in DOCCs (15, 37). We also tested *IR68a*, which is expressed in DOWCs for warming detection but is absent in DOCCs (17). Because of the difficulty in isolating DOCCs from the dorsal organ ganglion (DOG) (38), we collected the anterior region of larvae where DOCCs are located (Fig. S5*A*) and compared mRNA levels between control and *bishu-1^KO^* cells. At the early third instar stage, the mRNA levels of *IR25a* and *IR21a* were significantly reduced to approximately 50% of the control level in *bishu-1^KO^*, whereas *IR93a* and *IR68a* levels were not altered, suggesting that *bishu-1* could influence the expression of *Irs* in DOCCs (Fig. 5*A* and *B*). Contrarily, the levels of *IR25a* and *IR21a* did not differ between the control and *bishu-1^KO^* at the late third instar stage (Fig. S5*B*). When *IR25a* was overexpressed in *bishu-1^KO^* using *IR25a-GAL4* or *R11F02-GAL4*, the behavioral defect in discriminating between 20°C and 24°C was rescued (Fig. 5*C*). These results suggested that *bishu-1* is required to maintain the expression of cool temperature-sensitive *IR* genes and the responsiveness of DOCCs.

**Fig. 5.**
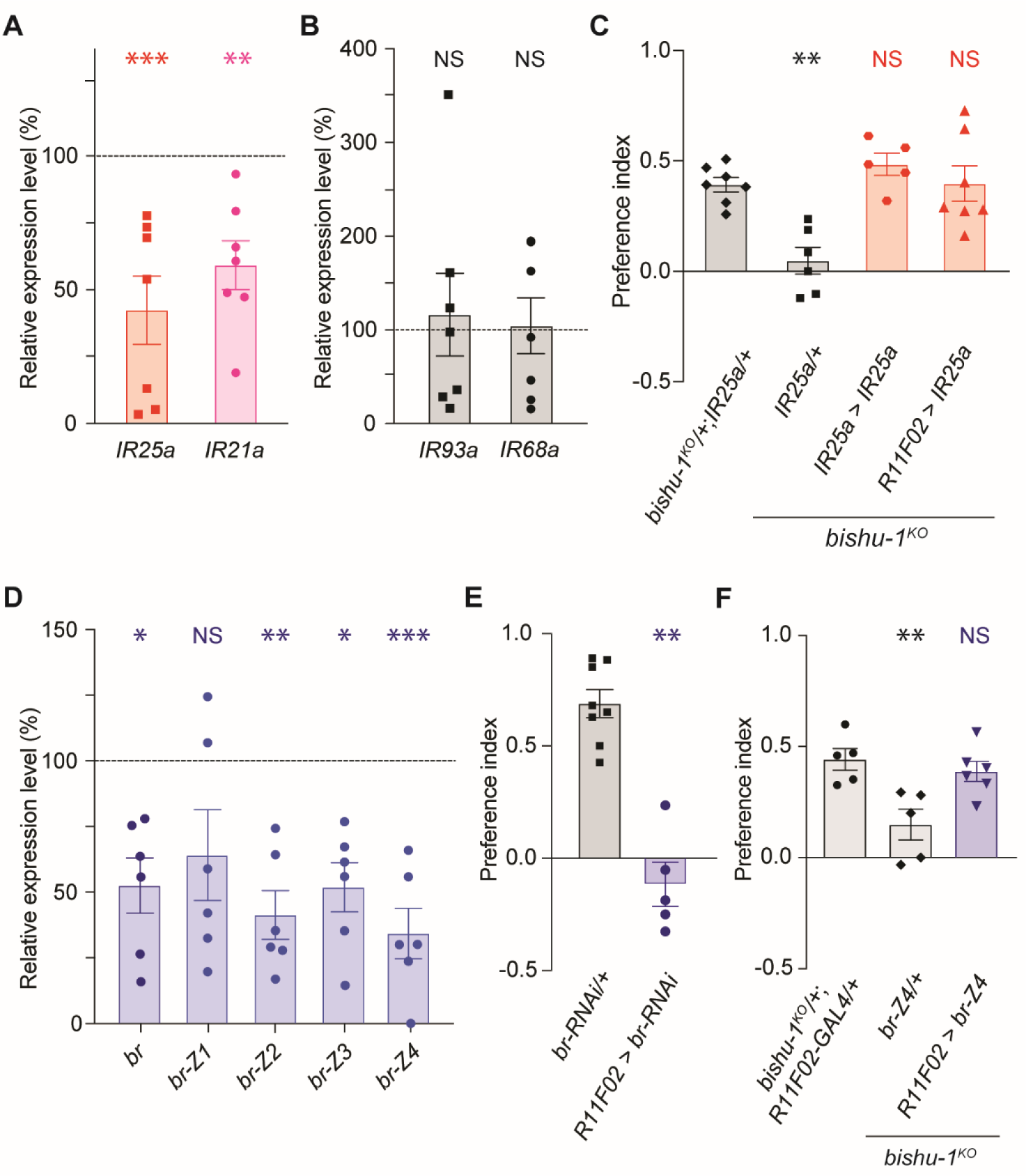
*bishu-1* regulated *IR25a* levels through the transcription factor *br*. (A and B) The relative expression of *IR25a* and *IR21a* (A); *IR93a* and *IR68a* (B) in *bishu-1^KO^* (N = 7). The expression of each gene was normalized to that of the control (*w^1118^*, 100%). (C) Rescue of *bishu-1^KO^* in two-way choice assays in the 20°C versus 24°C condition by overexpressing *IR25a* in *IR25a-*expressing neurons (*IR25a-GAL4*) or DOCCs (*R11F02-GAL4*, N = 5–7). (D) The relative expression of all isoforms (*br*) or a specific isoform (*br-Z1*, *Z2*, *Z3*, and *Z4*) in *bishu-1^KO^* (N = 6). The expression of each gene was normalized to that of the control. (E) The effect of *br* knockdown in DOCCs using *R11F02-GAL4* in the two-way choice assay in the 20°C versus 24°C condition (N = 5–8). (F) Rescue of *bishu-1^KO^* in the two-way choice assay in the 20°C versus 24°C condition by overexpressing *br-Z4* in DOCCs (*R11F02-GAL4*, N = 5–6). The data are presented as the mean ± SEM. \**P* < 0.05, \*\**P* < 0.01, and \*\*\**P* < 0.001 by one-way ANOVA with Dunnett’s test. NS denotes not significant.

To explore the possible interaction between *bishu-1* and *IR25a*, we sought transcription factors that are predicted to bind to the upstream region of *bishu-1* and *IR* genes. The database for transcription factor-binding sites (JASPAR 2022) suggested multiple candidates that are common or unique among *bishu-1* and *IR* genes, including six transcription factors upstream of the *IR25a* start codon (Fig. S5 *C* and *D*). Among these candidates, *broad* (*br*) expression was significantly reduced to approximately 50% of the control level, whereas other transcription factors were not suppressed (Fig. 5*D* and S5*E*). *br* has been reported to have at least four functional splicing variants (Fig. S5*F*) (39), and *br-Z2*, *br-Z3*, and *br-Z4*, but not *br-Z1*, were significantly suppressed in *bishu-1^KO^* (Fig. 5*D*). Consistent with this, knockdown of *br* in DOCCs using *R11F02-GAL4* resulted in no discrimination between 20°C and 24°C in larvae (Fig. 5*E*). Furthermore, the behavioral defect in *bishu-1^KO^* was rescued by overexpression of *br-Z4*, but not *br-Z1, br-Z2,* or *br-Z3* (Fig. 5*F* and S5*G*). Collectively, these results suggested that the *bishu-1*– dependent transcriptional regulation of *IR25a* through *br* contributes to the ability of DOCCs to regulate cooling temperature avoidance in larvae.

## Discussion

Our study provides evidence that the peripheral sensory process is maintained by the lipid metabolizing enzymes with putative MGAT activity, and one enzyme regulates the transcriptional level of the thermosensors *IR25a* and *IR21a* in DOCCs. Lipid metabolism is fundamental for many cellular functions ranging from cell membrane properties and lipid composition to intra-/intercellular signaling and energy storage (40, 41). In particular, DAG and its metabolites play key roles in regulating the activation of sensors and other membrane molecules, thereby affecting cellular and physiological responses (26, 42–44). In addition, a recent study clarified that *fa2h* that encodes a fatty acid 2-hydroxylase that maintains the cool temperature responsiveness of DOCCs in *Drosophila* larvae by regulating membrane rigidity through 2-OH sphingolipid production (21). Apart from these examples, our study revealed an unexpected link between DAG biosynthesis enzymes and the regulation of thermosensor expression, which contributes to the avoidance of cool temperatures in animals. *Drosophila* larvae exhibit a developmental switch in their thermal preference from warm (24°C) to cool temperature (18°C), and this switch depends on the rhodopsin signaling cascade in TRPA1-expressing neurons (4, 6) and Irs in DOCCs (37). The latter case involves the association of the thermal preference switch with the reduced expression of multiple *IR* genes at late larval stages (37). Whether the development-dependent regulation of *IR* expression is mediated by *bishu-1* requires further study.

We identified *br* as one of the transcription factors for the proper expression of *IR25a* (Fig. 5). The roles of *br* in metamorphosis and development have been well characterized (45), and multiple isoforms are expressed during central nervous system development (46). However, the regulatory mechanisms of *br* expression have not been clarified. Based on previous research and the current study, we can speculate several possible mechanistic links between *br* and *bishu-1*. For example, *br* is downregulated by harmful stimuli such as hypoxic stress (47) and toxic drug treatment (48). Because *bishu-1* contributes to lipid droplet formation through its MGAT activity (49, 50) and lipid droplets have been suggested to have a protective role against oxidative stress (51), it can be hypothesized that a reduction in lipid droplet content attributable to the absence of *bishu-1* increases oxidative stress, leading to a reduction in *br* expression. Another possible interaction between *bishu-1* and *br* might exist in some signaling cascades. In *Drosophila* larval brain neural stem cells, atypical PKC directly phosphorylates zinc-finger transcription factor (*zif*), thereby decreasing its activity by excluding it from the nucleus (52), whereas an interaction between *br* and *zif* was found in *Drosophila* transcription factor interactome analysis (53). Additionally, PKC supports the nuclear entry of *br* via RACK1 in silk moths (54). Because MGATs can derive the specific DAG isomer, 1,2-sn DAG, to activate PKCs (55, 56), *bishu-1*–dependent DAG production and PKC activation might control *br*–*zif* formation in DOCCs. Furthermore, *br* mediates the ecdysone signaling pathway, which regulates the dendritic pruning of *Drosophila* peripheral neurons during metamorphosis through AMPK-dependent gene regulation (57). Another study reported that *bishu-1* has an indirect interaction with the AMPK pathway in skeletal muscle performance (58). Therefore, it could be intriguing to examine the function of *bishu-1* and *br* in the AMPK signaling pathway during DOCC development.

We observed reduced responses of DOCCs to cooling stimulation in *bishu-1^KO^* (Fig. 4). In addition to the reduced expression of thermosensor *IR* genes (Fig. 5), the potential effects of neural development and maturation on the thermal response of DOCCs cannot be discounted. This possibility is raised by the fact that *IR25a* is required for the unique structural arrangement of the distal tips of the sensilla of cooling cells in adult *Drosophila* (16), which have corresponding functions as DOCCs in larvae. *Br* plays critical roles in neural development in the larval peripheral nervous system (46, 59), in addition to the transcription of *IR25a*. Thus, *br* could also be involved in the development of DOCCs to mediate the functional responses of *IR* genes. However, variation in dendritic bulb shape prevented us from comparing the control and *bishu-1^KO^*, and no clear structure changes were observed in the absence of *bishu-1* (Fig. S4). Conversely, we observed a shift in thermotaxis to cooler temperatures throughout larval development including the second and late third instar stages (Fig. S1), although we did not observe further reductions of *IR25a* and *IR21a* expression in *bishu-1^KO^* at the late third instar stage (Fig. S5). This result suggests that the abnormal cool temperature avoidance in *bishu-1^KO^* is not only attributable to changes in *IR* expression (17). Furthermore, a partial but not full rescue of the cooling response was observed upon overexpression of *bishu-1* in DOCCs. A previous report suggests that cross-inhibition exists between DOCCs and DOWCs in cooling/warming responses in DOG (17), where *bishu-1* and *IR25a* are co-expressed (Fig. 3). This provides a possibility that the cross-interaction mediates the temperature responses in *bishu-1*-expressing neurons.

*bishu-1* has been predicted to be a DGAT based on its sequence alignment and involvement in lipid droplet formation (49). However, given that the amino acid identity of *bishu-1* with human MGAT2 (39.52%) and MGAT3 (38.12%) are higher than that with human DGAT2 (30.67%), its molecular function as a DGAT has not been confirmed (49, 60). Our results demonstrated that multiple human MOGATs, but not DGAT2, compensated for the physiological function of *bishu-1* (Fig. 2), supporting the idea that *bishu-1* functions as a MGAT. As previously discussed, *bishu-1* possibly regulates the expression of *br* and *IR* genes through lipid droplet formation and/or DAG production. Further biochemical analyses will clarify the molecular function of *bishu-1* and the downstream mechanisms for gene regulation.

*bishu-1* forms a tandem structure together with *bisuh-2* and *CG1941* (Fig. S1). A previous study observed that while the transcriptional expression in the tandem is interdependent, these lipid enzymes with close molecular function displays independent roles (61). This support that although *bishu-1*, *bishu-2*, and *CG1941* display interactions at a mRNA level (Fig. S1), they contributed to thermotaxis distinctively. We identified *bishu-2* as a responsible gene for cool temperature avoidance (Fig. 1). The high amino acid identity between *bishu-1* and *bishu-2* (67.05%) suggests similar molecular functions. In contrast to *bishu-1*, *bishu-2* did not play a major role in DOCCs (Fig. 2) and functioned in other thermosensitive neurons such as *IR25a*-positive neurons (excluding *IR21a*-positive neurons) and chordotonal organs (Fig. S2). However, we haven’t identified a specific tissue where *bishu-2* plays the dominant role. We expect that *bishu-1* and *bishu-2* play distinctive and comprehensive roles in the thermotaxis of larvae. *CG1941* did not contribute to thermotaxis (Fig. S1), although it exhibited higher amino acid identity with *bishu-1* than *bishu-2* (69.60%). Comparing the expression pattern of *CG1941* to those of *bishu-1* and *bishu-2* could provide clues to understand their different physiological functions.

Taken together, our findings highlight the unpredicted role of *bishu-1* in cool temperature sensation through the regulation of thermosensors and a transcription factor. A number of investigations have revealed the function of *IR* genes in chemical sensation, thermal sensation, and hygrosensation (38, 62, 63), but little attention has been paid to their regulatory mechanisms. The current study provided novel insight into the functional correlation between lipid metabolism and sensory functions and proved the physiological importance of the coupling of lipid and sensors in appropriate sensory outputs.

## Materials and Method

### Fly stocks and husbandry

The following strains were obtained from the Bloomington *Drosophila* Stock Center (Bloomington, IN, USA): *bishu-1/2* transgenic BAC line (#90443), *UAS*-*dicer2* (#24650), *elav-GAL4* (#8760), *iav-GAL4* (#52273), *TRPL-GAL4* (#29134), *IR25a-GAL4* (#41728), *R11F02-GAL4* (#49828), *UAS-hMGAT2* (#82252), *UAS-hMGAT3* (#84925), *UAS-hDGAT2* (#84854), *40×UAS*-*IVS-mCD8GFP* (#32195), *10×QUAS-6×mCherry* (#52269), *5×UAS-mCD8::GFP* (#5137), *20×UAS-IVS-jGCaMP8m* (#92591), *UAS-IR25a* (#78067), *UAS-br-Z1* (#51190), *UAS-br-Z3* (#51192), and *UAS-br-Z4* (#51193). The following RNAi lines were obtained from the Vienna *Drosophila* Resource Center (Vienna, Austria): *UAS-bishu-1-RNAi* (#7942), *UAS-bishu-2-RNAi* (#108495), and *UAS-br-RNAi* (#104648). *IR21a-GAL4* was provided by P. Garrity. The following stocks were created in our laboratory: *CG1941^KO^*, *bishu-1^KO^*, *bishu-2^KO^*, *UAS-bishu-1*, *UAS-bishu-2*, *UAS-br-Z2*, and *bishu-1-P2AQF2*. We used *w^1118^* as the wild-type control, and all stocks used for behavioral assays were outcrossed with the control line for at least five generations. *UAS-dicer2* was combined with all RNAi lines to enhance the efficiency of knockdown.

Flies were reared on glucose–yeast–cornmeal medium [180 g of cornmeal, 100 g of dry brewer’s yeast Ebios, 19 g of agar, 250 g of glucose, 24 ml of methyl 4-hydroxybenzoate (10% w/v solution in 70% ethanol), and 8 ml of propionic acid in 2,500 ml of reverse osmosis (RO) water]. Flies were raised in vials or bottles in a 25°C incubator under a 12-h/12-h light/dark cycle.

### Phylogenetic comparison of MGATs and DGATs

The amino acid sequences of *Drosophila* and human proteins were obtained from GenBank, including *Drosophila melanogaster* CG1941 (NP_001286163), Bishu-1 (NP_610318), and Bishu-2 (NP_610319); human DGAT1 family members [diacylglycerol acyltransferase 1 (DGAT1, NP_036211), acetyl-CoA acetyltransferase 1 (ACAT1, NP_001373606), and ACAT2 (NP_001290182)]; and human DGAT2 family members [DGAT2 (NP_115953), DGAT2L6 (NP_940914), monoacylglycerol acyltransferase 1 (MGAT1, NP_477513), MGAT2 (NP_079374), MGAT3 (NP_835470), acyl-CoA wax alcohol acyltransferase 1 (AWAT1, NP_001013597), and AWAT2 (NP_001002254)]. The phylogenetic tree was constructed by the neighbor-joining algorithm implemented in the MEGA X program (64) using p-Distance (scale bar).

### Thermal preference assays

To prepare synchronized larvae for thermal gradient assays, 12–20 mated females were fed on yeast paste over 2 days and allowed to lay eggs in new food vials in a 3–6-h window. Larvae were raised in the vial until they reach the test stages (second instar, 48 h AEL; early third instar, 72 h AEL; late third instar, 120 h AEL). Larvae collected from the food were separated in 18% (72 and 120 h AEL) or 22% (48 h AEL) sucrose solution in 50-ml tubes (#1342-050S, Watson, Japan) twice, transferred to a 300 µm strainer (#43-50300-01, PluriSelect, Germany), followed by two rounds of washing with RO water. Larvae were kept in a 35 mm dish (#1000-035, Iwaki, Japan) at room temperature for 5–10 min to allow them to recover and subsequently used for the assays.

Thermal gradient assays were conducted following the previously described method (6, 20) with modifications. The test trays were assembled with an aluminum plate (22.6 × 6.0 × 0.1 cm^3^) and black acrylic wall (outer dimension: 20.0 × 6.0 cm^2^, inner dimension: 19.0 × 5.0 cm^2^, 0.5 cm height; Fig. 1*B* upper). The surface of the aluminum plate was covered with black aluminum tape (#J3270, Nitto, Japan). Trays were coated with 20 ml of 2% agarose. Two rods (18 × 0.5 × 0.5 cm^3^) were placed at the longer edges during agarose solidification to create grooves that minimize larva escape from the agarose after filling with water. The agarose surface was gently scratched and sprayed with water to prevent the gel from drying. Test trays were placed on top of two aluminum blocks separated by 18 cm and individually connected to a circulating water bath (NCB-1210A, Eyela, Japan) to generate a continuous linear temperature gradient from 8°C to 35°C (1.5°C/cm, Fig. 1*B* lower). The surface temperature of the agarose 0.5 cm from the wall was measured at every 2 cm with a digital thermometer (#MC3000, Chino, Japan).

To initiate the assay, larvae (40–65 individuals) were released in a line at 23°C (48 and 72 h AEL) or 29°C (120 h AEL). Each tray was covered with a square glass (20.0 × 6.0 × 0.1 cm^3^) with hydrophilic film (MF-600, Fujifilm Wako Pure Chemical, Japan) on both sides to prevent larval escape, heat loss, and fogging. Larvae at the second and early third instar stages were allowed to select temperatures on plates for 15 min, whereas tests were conducted for 10 min under LED red light (>600 nm) in a dark box for late third instar larvae. The distribution of larvae was captured by a CCD camera (#FL-CC1218-5MX, Ricoh, Japan) equipped with a 12-mm F1.8 Manual Iris C-Mount lens.

To quantify the distribution of larvae along the thermal gradient, the position of each larva in the image was determined by selecting the center of body using ImageJ (US National Institutes of Health, MD, USA) (65). The staying temperature of each larva was calculated using the following formula: staying temperature (°C) = horizontal distance from the position at 8°C (cm) × 1.5 (°C/cm) + 8°C. The number of larvae in each 2-cm zone (3°C range) was tabulated in the nine zones and divided by total number of larvae to calculate the proportion. We omitted larvae within 0.5 cm from both sides, those inside the water-filled grooves, those outside of the plates, and immobile larvae in the release zone from the calculation.

Thermal two-way choice assays were conducted on a test plate (outer: 14 × 10.1 × 0.9 cm^3^; inner: 12.9 × 8.7 × 0.8 cm^3^) coated with 25 ml of 2% agarose (Fig. *1F*). The plate was placed on top of two adjacent aluminum blocks that were separated using a plastic film (approximately 1 mm) as a spacer. The blocks were individually temperature-controlled using a circulating water bath. Agarose surfaces were gently scratched and sprayed with water to prevent gels from drying. The surface temperature on the center of each side of the test plate was measured and confirmed using a thermometer.

Early third instar larvae (72 h AEL, 40–65 larvae) were reared and collected as previously described in thermal gradient assays. They were released in a line at the border between two temperature zones and allowed to explore the tray under dim red LED light in a black acrylic box. After 15 min, the distribution of larvae on the plate was captured by a CCD camera, and the number of larvae in each temperature zone was tabulated (Fig. 1*F*). PI was calculated using the following formula: (number of larvae at 24°C – number of larvae at other temperatures)/(total number of larvae on the test tray). Larvae within the release zone (1 cm wide) were not counted in either temperature zone, and those outside the trays were not counted in the calculation.

### Confocal imaging

To examine the expression pattern of *bishu-1* in the DOG of larvae, we established *bishu-1* QF2 lines (*bishu-1-P2AQF2*) carrying *10×QUAS-6×mCherry* and *IR25-GAL4* or *IR21a-GAL4* carrying *40×UAS*-*IVS-mCD8::GFP*, and then we crossed the *bishu-1 QF2*-*mCherry* line with the *GAL4-GFP* line. The anterior region of larvae was dissected in phosphate-buffered saline (PBS) and mounted on a glass slide using 50% glycerol/PBS. GFP and mCherry fluorescence was captured using a confocal laser-scanning microscope (SP8, Leica, Germany) equipped with a HC PLAPO 63×/1.40 OIL CS2 objective lens. Z-sections were taken at 1-μm intervals. Images were analyzed using Leica Application Suite X (LAS X) and ImageJ software (65).

### Quantitative PCR

Whole-body sample (8–10 individuals) or dissected anterior regions from approximately 70 early third instar larvae (Fig. S5A) were homogenized in ice-cold PBS solution with a pestle. Total RNA was extracted with Sepasol-RNA I Super G (#0937984, Nacalai Tesque, Japan) and treated with Recombinant Dnase I (#2270A, Takara, Japan) for 30 min at 37°C, followed by denaturation with phenol:chloroform:isoamyl alcohol (25:24:1, #25970-14, Nacalai Tesque). RNA was precipitated with 100% ethanol and CH_3_COONa (#06893-24, Nacalai Tesque) by incubating at −20°C overnight. After obtaining total RNA, cDNA was synthesized from 2 mg of total RNA using ReverTra Ace qPCR RT Master Mix (#FSQ-201, Toyobo, Japan).

qPCR was performed using Thunderbird Next SYBR qPCR Mix (#QPX-201, Toyobo) and StepOne and QuantStudio3 Real Time PCR Systems (Applied Biosystems, MA, USA). qPCR primers are listed in Table S1. *Ribosomal protein 49* (*rp49*) was used as a reference gene. Gene expression was quantified with the ΔΔCt method, and the expression in mutants was normalized to that of a control sample.

### *In vivo* GcaMP-imaging

*in vivo* GcaMP imaging was conducted following a previously described method with modifications (11). To measure the calcium responses in DOCCs, *UAS-GcaMP8m* expression was driven by *R11F02-GAL4*. Early third instar larvae (72 h AEL) were prepared in the same manner as described for thermal preference assays. The anterior region of one larva was fixed on a 15-mm diameter disc made of silicone (KE-1606, Shin-Etsu, Japan) using insect pins (Austerlitz 0.1 mm, Czech Republic) in Ca^2+^-free HL3 solution contained 70 mM NaCl, 5 mM KCl, 10 mM NaHCO_3_, 20 mM MgCl_2_, 5 mM HEPES, 115 mM sucrose, and 5 mM trehalose (pH 7.2). DOCCs were imaged using a fluorescent microscope with a 25×/0.95 HC Fluotar water immerse lens (Leica). HL3 solution was perfused at 2.6 ml/min by a peristaltic pump (PSM071AA, Advantec, Japan). The temperature of the perfusate was controlled through an inline temperature controller (SC-20, Warner Instruments, MA, USA). The temperature near the larval sample was monitored using a temperature controller (CL-100, Warner Instruments) equipped with a thermistor probe (TA-29, Warner Instruments). Temperature fluctuation was recorded using AxoScope software (Molecular Devices, CA, USA). The GcaMP signal was captured every 3 s using a DFC9000 sCMOS camera (Leica) connected to the microscope (DM6 B, Leica) and recorded by LAS X (Leica). The fluorescence of images was processed with THUNDER in LAS X for computational clearing and StackReg in ImageJ for image stacks alignment, followed by subsequent analysis and measurement in ImageJ (65). Temperature fluctuations were analyzed using Clampfit 11.2 (Molecular Devices).

The Ca^2+^ responses in DOCCs were assessed by calculating changes in fluorescence intensity using the following formula: [(F_t_ – F_min_)/F_min_], where F_t_ and F_min_ represent the value obtained every 3 s and the minimum response during recording, respectively. The maximum response (F_max_) in each cooling phase was determined as the highest fluorescence intensity during 60–162 s (first) and 168– 270 s (second). The area under the curve in each cooling stage was calculated using a trapezoidal rule [(F_t_ + F_t +1_)/2 × 3 (sampling interval)] during 96–162 (first) and 204–270 s (second) and was normalized to the area under the curve of the first cooling phase in heterozygous controls.

### Statistical analysis

All data are presented as the mean ± standard error of mean (SEM) unless otherwise noted. The number of times each experiment was performed (*N*) is indicated in the Figure legends. The normality of distributions was assessed by the Shapiro–Wilk test. For pairwise comparisons, normally distributed samples were analyzed using an unpaired two-tailed Student’s *t*-test, and non-normally distributed samples were analyzed using the Mann–Whitney U test. For multiple pairwise comparisons, normally distributed samples were analyzed using one-way analysis of variance (ANOVA) with Dunnett’s or Tukey’s post-hoc test, and non-normally distributed samples were analyzed using the Kruskal–Wallis test with the Steel or Steel–Dwass test. Statistical analyses were performed using Prism 10 (GraphPad, CA, USA) or EZR (version 1.61; Saitama Medical Center, Jichi Medical University, Japan) (66), which is a graphical user interface for R (The R Foundation for Statistical Computing, Austria). Statistical significance was indicated by *P* < 0.05.

## Acknowledgments

The fly stocks used in this study were obtained from the Bloomington *Drosophila* Stock Center, and Vienna *Drosophila* Resource Center. We thank Dr. Paul Garrity (Brandeis University) for providing us fly stocks. We thank lab members Dr. Shoma Sato, Naomi Fukuta, Terumi Hashimoto, Aliyu Mudassir Magaji, and Hinata Yamanaka for supporting fly maintenance. We thank Dr. Shinji Takada (National Institute for Basic Biology, NIBB) and Dr. Minako Suzuki (NIBB) for providing the confocal microscopy Leica SP8 and supporting the imaging data collection. We thank Zhenbo Jiang (NIBB) for advising the transcription factor analysis using JASPAR 2022. This work was supported by Grant-in-aid for Scientific research 17H07337, 18K06495 and 21H02531 (for T. Sokabe) from Japan Society for the Promotion of Science (JSPS) and Ministry of Education, Culture, Sports, Science and Technology (MEXT) and Takeda Science Foundation (for T. Sokabe).

## Author Contributions

X.D. contributed to designing, conducting, and analyzing the experiments and preparing the draft and the final version of the manuscript. T. Sokabe contributed to designing and supervising the project and preparing the draft and the final version of the manuscript. T. Suito and M. Tominaga contributed to supervising the project and preparing the final version of the manuscript.

