## Supplemental information for "Monoacylglycerol acyltransferase maintains ionotropic receptor expression for cool temperature sensing and avoidance in *Drosophila*"

### Extended Methods

#### Generation of knockout, transgenic, and reporter flies

*CG1941$^{KO}$*, *bishu-1$^{KO}$*, and *bishu-2$^{KO}$* fly lines were generated using the CRISPR-Cas9 system. *CG1941$^{KO}$* was generated by non-homologous end joining with a 5-bp deletion at +55–59 bp (the start codon of *CG1941* was designated as position 1, Fig. S1*B*, top). The guide RNA (gRNA) sequence (5'-GCGTCGCCAGACGTTTGCCA-3') was cloned into the pU19_DsRed_U6_3 vector (C. Montell) and microinjected into embryos carrying *vas-cas9* on X chromosome (BL #55821) by BestGene (CA, USA). The transformants were crossed with *w$^{1118}$*, and the progeny were screened for gene targeting by DNA sequencing. *bishu-1$^{KO}$* and *bishu-2$^{KO}$* were generated by homology-directed repair (HDR) with *DsRed* insertion at +33 and +28 bp, respectively (Fig. S1*B*, middle and bottom). The gRNA sequence for *bishu-1* (5'-ATCTGCAGTCGGCGTTCCAG-3') or *bishu-2* (5'-GGCTCCTCTCCGGGTTCCGC-3') was cloned into the pU19_DsRed_U6_3 vector. Approximately 1-kb fragments upstream and downstream of the Cas9-mediated double-strand break sites were PCR-amplified and subcloned into the pHD-Scarless-DsRed donor vector (#64703, Addgene, MA, USA) using the NEBuilder HiFi DNA Assembly kit (NEB, MA, USA). The gRNA vector and donor vector were injected into *vas-cas9* (X) embryos by BestGene. The wild-type sequences of neighboring genes were confirmed in each mutant using primers for donor vector construction (*bishu-1* and *bishu-2* in *CG1941$^{KO}$*, *CG1941* and *bishu-2* in *bishu-1$^{KO}$*, and *CG1941* and *bishu-1* in *bishu-2$^{KO}$*). PCR primers for vector constructions, genotyping, and sequence confirmation are listed in Table S1.

*UAS-bishu-1*, *UAS-bishu-2*, and *UAS-br-Z2* lines were generated using the PhiC31 site-specific integration system. The PCR fragment of the *bishu-1* coding sequence was cut with *Xho*I/*Xba*I and inserted into the *Xho*I/*Xba*I site of the pJFRC7-20×UAS-IVS-mCD8::GFP vector (#26220, Addgene). The PCR fragment of the *bishu-2* coding sequence was cut with *Xho*I/*Spe*I and inserted into the *Xho*I/*Xba*I site of the vector. The PCR fragment of *br* Z2 isoform were inserted into the vector using NEBuilder HiFi DNA Assembly kit (NEB). The *bishu-1*, *bishu-2*, or *br-Z2* vector was microinjected into P{CaryP}55C4 embryos (BL #8622) by BestGene. PCR primers for vector constructions and genotyping are listed in Table S1.*bishu-1-P2AQF2* was generated by replacing the stop codon of the gene with *QF2* and the *mini-white* marker (Fig. S3*A*). The gRNA sequence (5'-AAACTAGTGTACAACTAGAG-3') was cloned into the pU6-3_BbsI_chiRNA vector (#45946, Addgene). An approximately 1-kb fragment upstream of the *bishu-1* stop codon was PCR-amplified and subcloned into the *Kpn*I site of the pW35P2AQF2 donor vector (C. Montell). The NEBuilder HiFi DNA Assembly kit was used to subclone the approximately 1-kb downstream fragment. Both the gRNA vector and donor vector were microinjected into *vas-cas9* (X) by BestGene. PCR primers for vector constructions and genotyping are listed in Table S1.

#### Measurement of developmental rates

The timings of third instar entry and pupation were determined following the previously described method (1). Flies were prepared for egg laying in the same manner as for the temperature gradient assay. Fifteen larvae were collected 74 h AEL and transferred to a glass slide, and their mouth hooks and spiracles were examined under a Nikon TE300 microscope using a 40 × /0.55 NA Ph1 ADL lens (#MRP46402, Nikon) to determine the proportion of third instar larvae. Larvae at the transition stage from second to third instar displaying two pairs of mouth hooks (2) were treated as third instar larvae. The pupation timing was assessed by counting the number of pupae on the vial walls every 12 h during light periods (ZT0 and ZT12) starting from 108 h AEL. The percentages of pupae were calculated according to the maximum number at 228 h AEL. T50 was the time at which 50% of the animals underwent pupation.

**Confocal imaging**

To examine the morphology of DOCCs in *bishu-1*$^{KO}$ larvae, we established the *R11F02-GAL4* line carrying *5×UAS-mCD8::GFP* under a *bishu-1*$^{KO}$/+ or *bishu-1*$^{KO}$ background. The anterior region of larvae was dissected in PBS and mounted on a glass slide using 50% glycerol/PBS. GFP fluorescence was captured using a confocal laser-scanning microscope (FV1200, Olympus, Japan) equipped with a UPL SAPO 60 × S objective lens. Z-sections were taken at 0.5-μm intervals. Images were analyzed using FLUOVIEW (version 4.2c, Olympus) and ImageJ software (3).

**Table S1.** Primers used in this study.

| **Name** | **Primers (5′ to 3′)** |
|---|---|
| Primers for generating knockout, knock-in, and transgenic lines | |
| *CG1941_gRNA_S* | ACGTTTGCCAGTTTTAGAGCTAGAAATAGC |
| *CG1941_gRNA_AS* | CTGGCGACGCCGACGTTAAATTGAAAATAGG |
| *bishu-1_gRNA_S* | GGCGTTCCAGGTTTTAGAGCTAGAAATAGC |
| *bishu-1_gRNA_AS* | GACTGCAGATCCGACGTTAAATTGAAAATAGG |
| *bishu-2_gRNA_S* | CGGGTTCCGCGTTTTAGAGCTAGAAATAGC |
| *bishu-2_gRNA_AS* | GAGAGGAGCCCGACGTTAAATTGAAAATAGG |
| *HDR_Arm1-F* | CGTTTCACTTCTGAGTTCGG |
| *HDR_Arm1-R* | CTCTTATACGACATCACCGATG |
| *HDR_Arm2-F* | GCAGTATACGAGACCTATAGG |
| *HDR_Arm2-R* | GTATAGGAGACCTATAGTGTCTTC |
| *bishu-1_Arm1_F* | CCATCGGTGATGTCGTATAGGAATTGGCGCTCCCATCGACGTG |
| *bishu-1_Arm1_R* | CCTATAGGTCTCGTATACTGCCAGAGGAACCCGCAGTGGTGC |
| *bishu-1_Arm2_F* | GAAGACACTATAGGTCTCCTATACGAACGCCGACTGCAGATACTGG |
| *bishu-1_Arm2_R* | CATCGGTGATGTCGTATAAGAGACAGTCCTGAAACCGTCGGAGC |
| *bishu-2_Arm1_F* | CCATCGGTGATGTCGTATAGGAAGCTTCCTCCCATTCCGACGACG |
| *bishu-2_Arm1_R* | CCTATAGGTCTCGTATACTGCGAACCCGGAGAGGAGCCCATTC |
| *bishu-2_Arm2_F* | GAAGACACTATAGGTCTCCTATACCGCTGGAACGGCGGCTTC |
| *bishu-2_Arm2_R* | CATCGGTGATGTCGTATAAGAGACCATGGCTTCCTTGGCGCCAC |
| *bishu-1_UAS_F* | AAAACTCGAGATGAAAATCGAGTGGGCACCAC |
| *bishu-1_UAS_R* | TTCTAGACTAGTGTACAACTAGAGTGGCAC |
| *bishu-2_UAS_F* | AAAACTCGAGATGACAATCGAATGGGCTC |
| *bishu-2_UAS_R* | TTCTAGATCACTGTATTATAAGTTTGATATGC |
| *br-Z2_UAS-F* | TCAGGCGGCCGCGGCTCGAGATGGACGACACACAGCACTTC |
| *br-Z2_UAS-R* | CTTCACAAAGATCCTCTAGATCAGATCGACGAGTTGAACTGG |
| *bishu-1_knockin_gRNA_S* | ACAACTAGAGGTTTTAGAGCTAGAAATAGC |
| *bishu-1_knockin_gRNA_AS* | ACACTAGTTTCCGACGTTAAATTGAAAATAGG |
| *bishu-1_knockin_Arm1_F* | AAGGTACCTTGGCCTACGTCTTTGTGC |
| *bishu-1_knockin_Arm1_R* | AAGGTACCGTGTACAACTAGAGTTGCACTCTTC |
| *bishu-1_knockin_Arm2_F1* | ATACATACTAGGCGCGCCCAGGCCGCTAGTTGTACACTAG |
| *bishu-1_knockin_Arm2_F2* | TGGGCGCGCCTAGTATGTATGTAAG |
| *bishu-1_knockin_Arm2_R1* | AGCTGAGCAAACAAACAAGCGCCCAGCAGGATAGCTACTG |
| *bishu-1_knockin_Arm2_R2* | GCTTGTTTGTTTGCTCAGCTTACG |
| Primers for genotyping | |
| *CG1941*$^{KO}$ forward | GAGGACCGTGCTACCAAGTAG |
| *CG1941*$^{KO}$ reverse | TGCGGTCTTGACCAGCTGTAC |
| *bishu-1*$^{KO}$ forward | ATCGCTGGCTTCCATATCATTC |
| *bishu-1*$^{KO}$ reverse | CCAGCGACTCAATGACCTGT |
| *bishu-2*$^{KO}$ forward | ATAAGTACTCCACGTTACACCCTGG |
| *bishu-2*$^{KO}$ reverse | TCTCCAATTGCTGCGGTAGAA |
| *vector_UAS_g_F* | GCAGCTGAACAAGCTAAACAATC |
| *bishu-1_UAS_g_R* | TGCCATCCACAACGGATTG |
| *bishu-2_UAS_g_R* | CTCCGATTGCTGCGGTAGAAT |
| *br-Z2_UAS_g_R* | AAGGACTGCAGGGACTTCTGGTG |
| *bishu-1-P2AQF2* forward | CACCTTTGGCTTCCTCCCAT |
| *bishu-1-P2AQF2* reverse | GAGCGTTAGCCTCAGCCGCAG |
| Primers for qPCR | |
| *rp49* forward | GACGCTTCAAGGGACAGTATCTG |
| *rp49* reverse | AAACGCGGTTCTGCATGAG |

| | |
|---|---|
| *CG1941* forward | TGCGGTCTTGACCAGCTGTAC |
| *CG1941* reverse | CTATTTCTTCGTTGCTGCCGTG |
| *bishu-1* forward | CACCTTTGGCTTCCTCCCAT |
| *bishu-1* reverse | AGTGGCACTCTTCGAATTCTCC |
| *bishu-2* forward | TGCCTCGCAGTAGCTATCCT |
| *bishu-2* reverse | CTCCGATTGCTGCGGTAGAAT |
| *IR25a* forward | AGTCAGCGGGACAATGCGAC |
| *IR25a* reverse | CGTGACGAGATCAAAGGTTCCATAC |
| *IR93a* forward | TCTAAATTGGAAGACCGCCGTTG |
| *IR93a* reverse | GTTCAGGGTCTCGGCTATTTCGATG |
| *IR68a* forward | ATTGCCATCAGTCGGTATCGTTC |
| *IR68a* reverse | ATCATCGCTGTAGCCGAACAC |
| *IR21a* forward | CTCAATAAATGCCCACCGGTC |
| *IR21a* reverse | TGCAATTGCAATCTATATGGCTCG |
| *broad* common forward | CTACTTCCGCGAGCTGCTCAAG |
| *broad* common reverse | AAGGACTGCAGGGACTTCTGGTG |
| *br*-isoform common forward | TATCGATCCCTCAGCGTTGCTAG |
| *br-Z1* reverse | ATTGACAGGCCACCGTTGCTC |
| *br-Z2* reverse | GCAGAGGAGCTTACCGCAGAG |
| *br-Z3* reverse | GGTGGCGATGAGGTAGAACTG |
| *br-Z4* reverse | GGTGTGGTGATGACTGTTGTTCTAC |
| *GATA-d* forward | AACTACGGCAAAGAGCACGGTC |
| *GATA-d* reverse | CCACAAACCACGTTGGGTAATC |
| *pan* forward | ATGAGCACGGAAGTCAGCTAAG |
| *pan* reverse | CAATGTGGGTCGTTGCTGTG |
| *Abd-B* forward | CCACTGCATATACCCGCCAT |
| *Abd-B* reverse | TCCGCTTCGTTCATGTAGGC |
| *Dr* forward | ATGTTTCCGGGAGCAGGATTC |
| *Dr* reverse | AGCAGCTGCTGTGTTGTGAAG |
| *exd* forward | GCAACCCATATCCATCCGAAGAG |
| *exd* reverse | TGGATAGCCCATGGAATCCTG |
| *Ubx* forward | GAGTCCCTATGCCAACCACC |
| *Ubx* reverse | AGGCAGTCCTGTTTGTAGGC |
| *Vsx2* forward | CGAAATGCTCTCGCTGAAGAC |
| *Vsx2* reverse | CCTTGGCCGACTTAAGGATCGTG |

Fig.S1

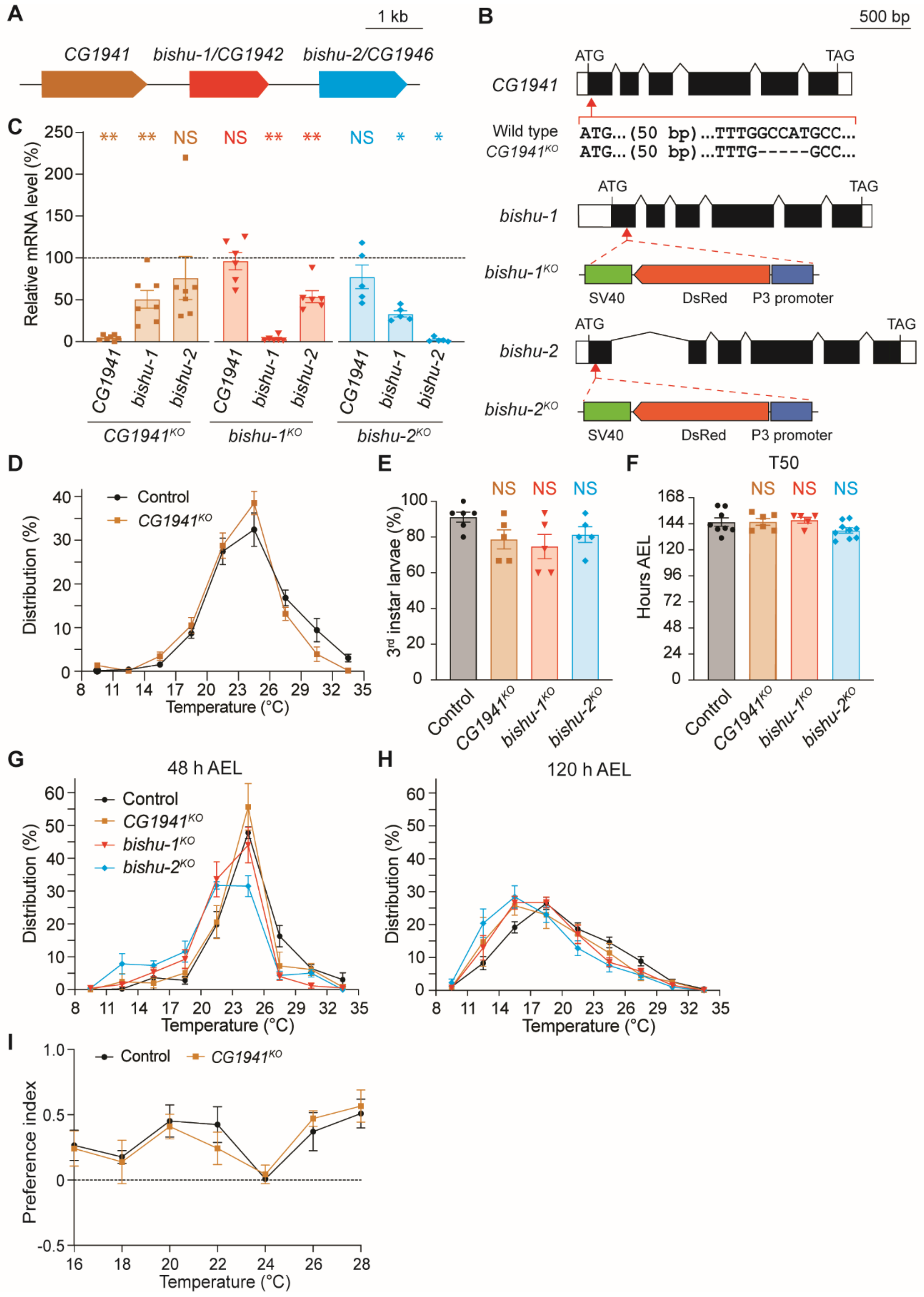

A
1 kb
CG1941
bishu-1/CG1942
bishu-2/CG1946
B
500 bp
ATG
TAG
CG1941
Wild type
CG1941KO
ATG...(50 bp)...TTTGGCCATGCC...
ATG...(50 bp)...TTTG-----GCC...
bishu-1
bishu-1KO
SV40
DsRed
P3 promoter
bishu-2
bishu-2KO
C
Relative mRNA level (%)
CG1941KO
bishu-1KO
bishu-2KO
D
Distribution (%)
Control
Temperature (°C)
E
3rd instar larvae (%)
F
T50
Hours AEL
G
48 h AEL
H
120 h AEL
I
Preference index

**Fig. S1. Mutant design, thermal preferences and developmental rates of *CG1941*, *bishu-1*, and *bishu-2*.**

(A) The relative position of *CG1941* (brown), *bishu-1* (red), and *bishu-2* (light blue) on the genome.

(B) Gene structures and mutation designs of $CG1941^{KO}$ (top), $bishu\text{-}1^{KO}$ (middle), and $bishu\text{-}2^{KO}$ (bottom). White and black boxes indicate untranslated regions and exons, respectively. ATG and TAG indicate start and stop codons, respectively. Spaces between exons indicate introns. Mutants were generated using the CRISPR-Cas9 technique. $CG1941^{KO}$ (top) contains a 5-bp deletion at 55 bp downstream of the start codon after non-homologous end joining, which causes a frame shift. $bishu\text{-}1^{KO}$ (middle) and $bishu\text{-}2^{KO}$ (bottom) feature a *DsRed* insertion in the first exon.

(C) Relative mRNA levels of $CG1941^{KO}$, $bishu\text{-}1^{KO}$, and $bishu\text{-}2^{KO}$ in the whole bodies of early third instar larvae (72 h AEL). mRNA levels in the mutants were normalized to that of the control ($w^{1118}$, 100%, N = 5–7).

(D) Distribution of early third instar larvae of the control and $CG1941^{KO}$ on an 8°C–35°C linear gradient (N = 6–7).

(E) The percentage of larvae entering the third instar stage at 74 h AEL in the control, $CG1941^{KO}$, $bishu\text{-}1^{KO}$, and $bishu\text{-}2^{KO}$ (N = 5–6).

(F) The timing of pupation required for 50% of larvae to become pupae in the control, $CG1941^{KO}$, $bishu\text{-}1^{KO}$, and $bishu\text{-}2^{KO}$ (N = 5–8).

(G and H) Distribution of second (48 h AEL, F) and late third instar larvae (120 h AEL, G) in the control, $CG1941^{KO}$, $bishu\text{-}1^{KO}$, and $bishu\text{-}2^{KO}$ on an 8°C–35°C gradient (N = 7–8).

(I) Preference indices of control, $CG1941^{KO}$, $bishu\text{-}1^{KO}$, and $bishu\text{-}2^{KO}$ larvae between 24°C and other temperatures (16°C, 18°C, 20°C, 22°C, 24°C, 26°C, and 28°C) in the thermal two-way choice assay (N = 3–6).

The data are presented as the mean ± SEM. *$P$ < 0.05 or **$P$ < 0.01 by the Kruskal–Wallis test with Steel's multiple comparison versus the control (C). NS denotes not significant. In (E and F), No significant differences were observed by the one-way ANOVA with Dunnett's test.

Fig. S2

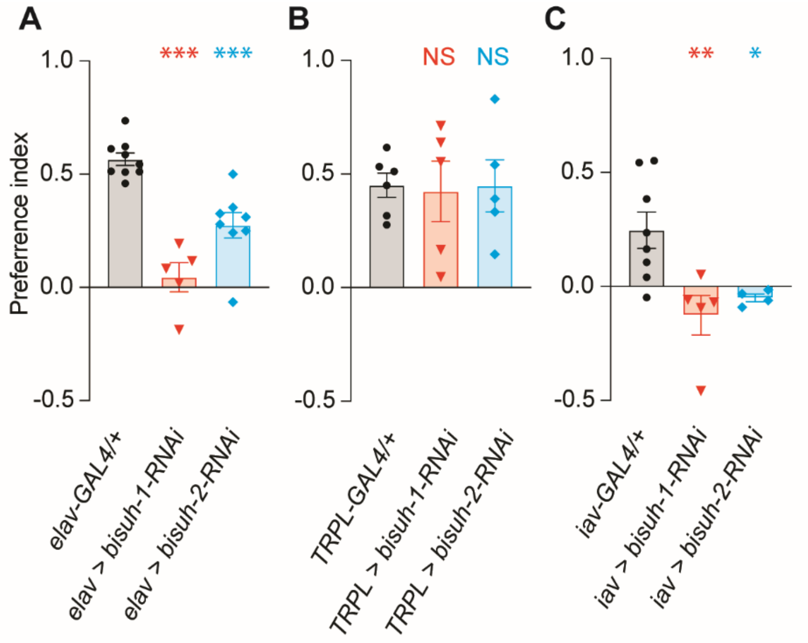


**Fig. S2. Contribution of *bishu-1* and *bishu-2* in neurons to cool temperature avoidance.**

(A–C) The effects of *bishu-1* or *bishu-2* knockdown on the Preference indices of the thermal two-way choice assay in a 20°C versus 24°C condition with the indicated *GAL4* lines: *dicer-2*;*elav-GAL4* (A, N = 5–9), *dicer-2*;*TRPL-GAL4* (B, N = 5–6), and *dicer-2*;*iav-GAL4* (C, N = 4–8). The data are presented as the mean ± SEM. **P* < 0.05, ***P* < 0.01, and ****P* < 0.001 by one-way ANOVA with Dunnett's multiple comparison test. NS denotes not significant.

Fig.S3

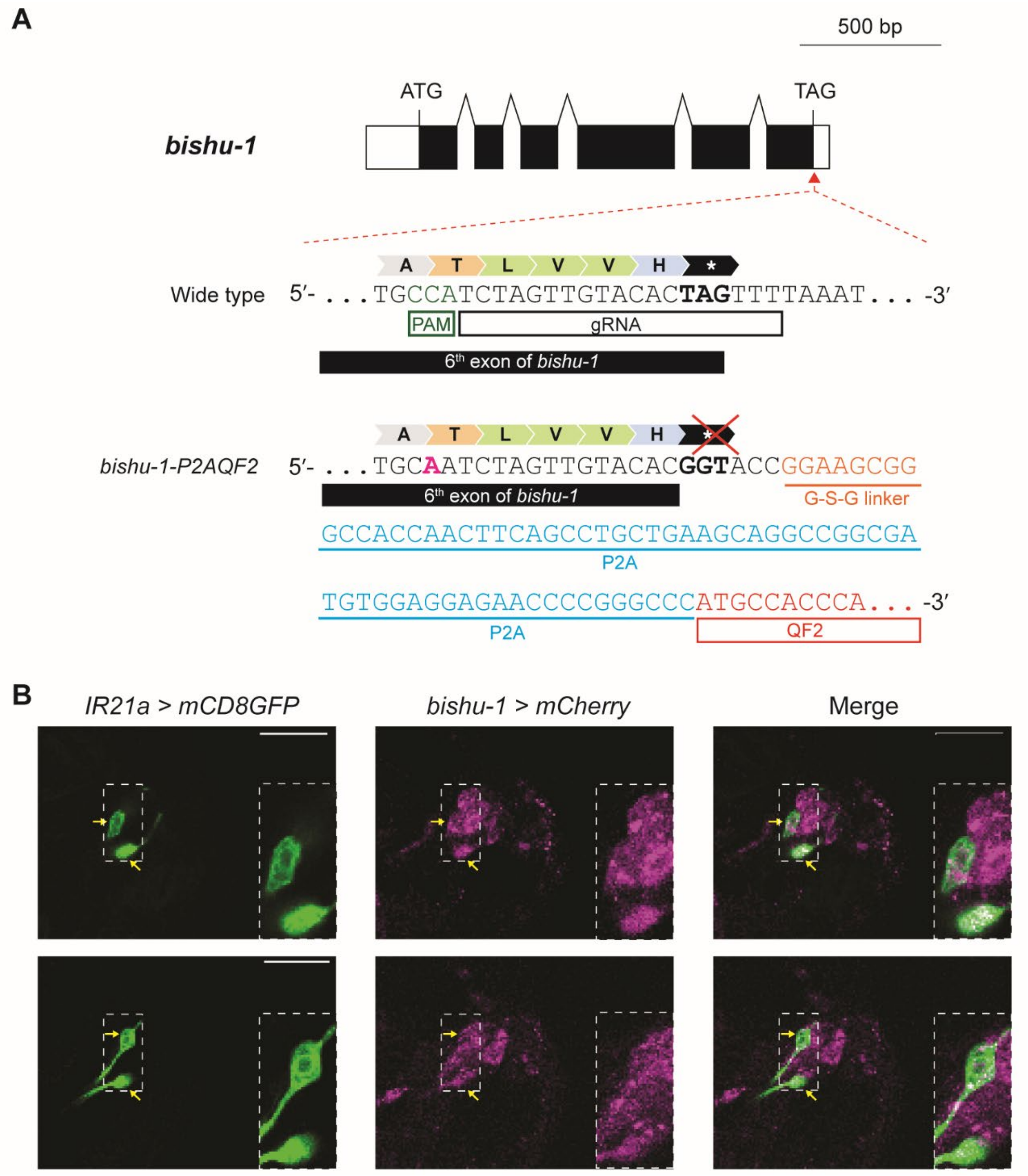
A
500 bp
ATG
TAG
bishu-1
A T L V V H *
Wide type 5′- . . . TGCCATCTAGTTGTACACTAGTTTTAAAT . . . -3′
PAM
gRNA
6th exon of bishu-1
A T L V V H *
bishu-1-P2AQF2 5′- . . . TGCAATCTAGTTGTACACGGTACCGGAAGCGG
6th exon of bishu-1
G-S-G linker
GCCACCAACTTCAGCCTGCTGAAGCAGGCCGGCGA
P2A
TGTGGAGGAGAACCCCGGGCCCATGCCACCCA . . . -3′
P2A
QF2
B
IR21a > mCD8GFP
bishu-1 > mCherry
Merge

**Fig. S3. The design and expression pattern of *bishu-1-P2AQF2* in DOCCs.**
(A) Design of gRNA on wild-type *bishu-1* and the insertion of *P2AQF2* downstream of *bishu-1*. The Cas9-targeted protospacer adjacent motif (PAM) used in this modification is indicated (green). To avoid the cleavage of homologous Arm in *bishu-1-P2AQF2*, a synonymous substitution was introduced in the corresponding region ('C' to "A', pink). G-S-G linker, P2A sequence, QF2 coding region, and *mini-white* marker (not shown) were inserted immediately upstream of the stop codon.
(B) Single optical sections of *IR21a-GAL4* (left, *IR21a-GAL4/*+;*UAS-40×GFP/*+), *bishu-1-P2AQF2* (middle, *10×QUAS-6×mCherry/*+,*bishu-1-P2AQF2/*+), and the merged image (right) from Figure 3, D–F. The positions of the cell bodies of DOCCs are indicated by arrows and highlighted with magnified insets from two identical slices (upper and lower). In all images, the right is the anterior side. Scale bars represent 20 μm.

Fig.S4

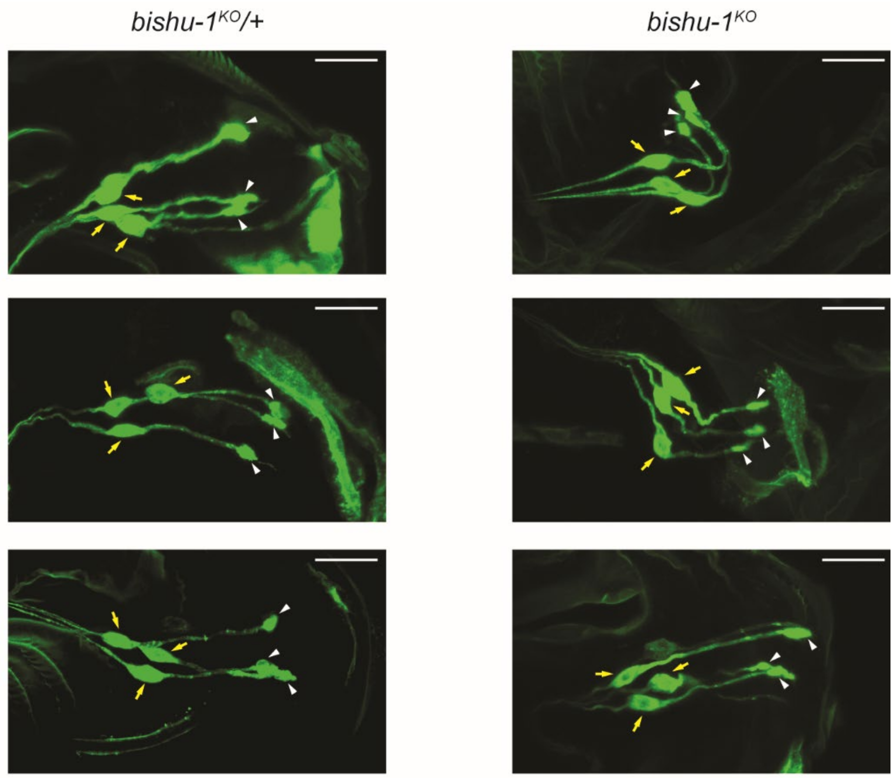


**Fig. S4. The morphology of DOCCs in the control and *bishu-1$^{KO}$.***
Three representative images of the morphology of GFP-expressing DOCCs in the control (left, *5×UAS-GFP*,*bishu-1$^{KO}$/*+;*R11F02-GAL4/*+) and *bishu-1$^{KO}$* (right, *5×UAS-GFP*,*bishu-1$^{KO}$/bishu-1$^{KO}$*;*R11F02-GAL4/*+). Arrows and arrowheads indicate the cell bodies and dendritic bulbs of DOCCs, respectively. The expression pattern was investigated in more than 15 individuals). In all images, the right is the anterior side. Scale bars represent 20 μm.

Fig.S5

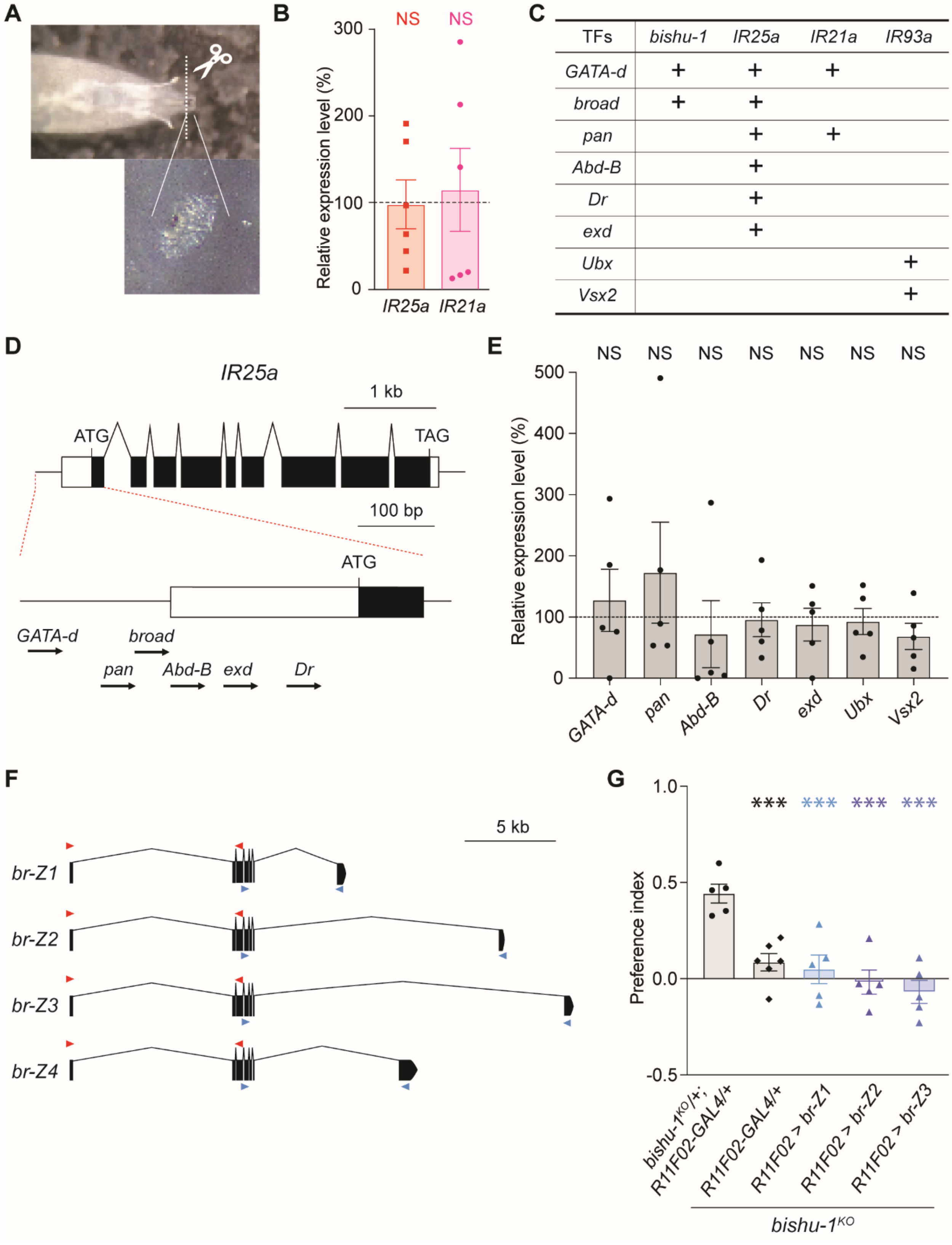
A
B
NS
NS
300
200
100
0
Relative expression level (%)
IR25a
IR21a
C
TFs
bishu-1
IR25a
IR21a
IR93a
GATA-d
broad
pan
Abd-B
Dr
exd
Ubx
Vsx2
D
IR25a
1 kb
ATG
TAG
100 bp
ATG
GATA-d
broad
pan
Abd-B
exd
Dr
E
NS
500
400
300
200
100
0
Relative expression level (%)
GATA-d
pan
Abd-B
Dr
exd
Ubx
Vsx2
F
5 kb
br-Z1
br-Z2
br-Z3
br-Z4
G
1.0
0.5
0.0
-0.5
Preference index
***
bishu-1KO/+; R11F02-GAL4/+
R11F02-GAL4/+
R11F02 > br-Z1
R11F02 > br-Z2
R11F02 > br-Z3
bishu-1KO

**Fig. S5. The expression of transcription factor candidates that bind upstream of *bishu-1* and *IR* genes and their involvement in thermal responses.**
(A) A representative image presenting part of the cut anterior region of a larva for collecting thermosensory neurons including DOCCs. Anterior regions from approximately 70 individuals in each genotype were collected and processed for RNA extraction.
(B) The relative expression of *IR25a* and *IR21a* in *bishu-1*$^{KO}$ at the late third instar stage (120 h AEL, N = 6). The expression of each gene was normalized to that of the control at the same stage.
(C) Existence (+) of predictive binding sites of transcription factors upstream of the *bishu-1*, *IR25a*, *IR21a*, and *IR93a* coding regions. The binding was predicted by TFBS predictions in the *Drosophila melanogaster* genome (genome: dm6) using the JASPAR CORE insect collection (https://jaspar2020.genereg.net/matrix-clusters/insects/).
(D) Gene structure of *IR25a* (upper) and positions of predictive transcription factor-binding motifs (lower). Arrows indicate the direction of transcription for each factor.
(E) The relative expression of *GATA-d*, *pan*, *Abd-B*, *Dr*, *exd*, *Ubx*, and *Vsx2* in *bishu-1*$^{KO}$ at the early third instar stage (72 h AEL, N = 5). The expression of these genes was normalized to that of the control.
(F) The structure of four functional variants of *br*. Arrowheads indicate the position of qPCR primers that cover common (red) and isoform-specific regions (blue).
(G) Preference indices of the thermal two-way choice assay in a 20°C versus 24°C condition at the early third instar stage (72 h AEL). Overexpression of *br* isoforms (*br-Z1*, *br-Z2*, and *br-Z3*) in *bishu-1*$^{KO}$ using the DOCC-specific driver *R11F02-GAL4* (N = 5–6). The data are presented as the mean ± SEM. ****P* < 0.001 by one-way ANOVA with Dunnett's test. NS denotes not significant.
